# ARPE-19 Epithelial Cells Fail To Initiate Type-I Interferon Signaling in Response to Human Cytomegalovirus Infection

**DOI:** 10.1101/2024.08.01.606090

**Authors:** Mariana Andrade-Medina, Todd M. Greco, Ileana M. Cristea, Adam Oberstein

**Affiliations:** University of Illinois at Chicago, Department of Microbiology and Immunology; Princeton University, Department to Molecular Biology

## Abstract

ARPE-19 cells are a commonly used epithelial model for studying human cytomegalovirus (HCMV) infection. We recently found that ARPE-19 cells assume a mesenchymal phenotype when maintained at low confluency and that ARPE-19 cells resemble mesenchymal fibroblasts rather than epithelial cells in HCMV infection assays. Here, using comparative proteomics analysis, we find that subconfluent ARPE-19 cells are also deficient in their ability to initiate canonical type-I interferon signaling. Comparative proteomic analysis between subconfluent ARPE-19 and MRC-5 cells revealed a lack of canonical type-I interferon response in ARPE-19 cells upon HCMV infection, evidenced by the absence of interferon stimulated gene (ISG) induction. qRT-PCR and RNA-sequencing analysis revealed that ARPE-19 cells fail to initiate *interferon-beta* transcription in response to HCMV infection, yet they are competent to respond to exogenously interferon-b, indicating a failure in early pathogen detection. ARPE-19 cells showed low baseline levels of key intracellular pattern recognition receptors (PRRs) such as CGAS and IFI16, as well as the signaling molecule STING. This deficiency was associated with a failure to activate IRF3 phosphorylation, a crucial step in interferon signaling. These findings suggest an upstream defect in the early detection of viral components, likely due to reduced expression of critical PRRs. ARPE-19 cells may be inherently deficient in initiating interferon responses due to their derivation or possibly due to their origin from an immune-privileged tissue. Our results continue to highlight important phenotypic characteristics of the ARPE-19 cell line; important considerations for those using ARPE-19 cells as an experimental infection model for studying HCMV or other human viruses.

## Introduction

The ARPE-19 cell line is a commonly used retinal epithelial cell line for biochemical and virological studies^1–19^. Despite being derived from a retinal epithelium explant^20^, ARPE-19 cells exhibit cell culture characteristics that diverge significantly from primary retinal epithelial cells^21–25^. Most importantly, ARPE-19 cells exhibit strong density-dependent changes in their epithelial phenotype^20,21,26,27^. At low cell density, typical of subcultured, or “passaged” cells in monolayer cell culture, ARPE-19 cells exhibit a mesenchymal morphology and express low levels of genes involved in epithelial functional and retinal biogenesis^21,27^. A number of studies have found that long periods of high confluency culture are required for ARPE-19 cells to establish a functional epithelial phenotype^20,21,23,27–29^. After two to four months of high density culture, ARPE-19 cells turn on expression of epithelial (e.g., CDH1 and EPCAM) and visual cycle (e.g., RPE1, LRAT, RDH5, and RDH10) genes^21,23,27^; assume a cobblestone epithelial morphology^20,23,27^; establish apico-basal polarization^20,28,29^; and display high trans-epithelial resistance (a measure of barrier function)^20,27–29^.

Despite their known density-dependent phenotype, ARPE-19 cells are frequently used as a simple epithelial cell culture model for experimental infection studies with a variety of human viruses ^30–37^, including human cytomegalovirus (HCMV)^1,3–8,10–13,15–18,21,38–40^. We recently found that subconfluent (i.e. “just confluent”) ARPE-19 cells resemble mesenchymal fibroblasts, rather than epithelial cells, in experimental HCMV infection assays^21^. Gene expression profiling experiments showed that ARPE-19 cells in a low confluency state mirrored fibroblasts, contrastingly markedly with other epithelial cell lines with strong intrinsic epithelial character^21^. For example, subconfluent ARPE-19 cells were permissive for all phases of the HCMV infectious cycle, producing logarithmic increases of cell-free infectious progeny; whereas strongly epithelial cell lines, such as MCF10A or RWPE-1, restricted HCMV biosynthesis and failed to yield significant amounts of cell-free infectious progeny^21^.

Here, by comparing the cellular proteomic response to infection across ARPE-19 and MRC-5 cells, we find that subconfluent ARPE-19 cells fail to initiate a canonical type-I interferon response to HCMV infection. Infected ARPE-19 cells fail to induce interferon beta (*IFNB*) transcription in response to HCMV infection, as well as transcription and translation of downstream interferon stimulated genes (ISGs). Treatment of ARPE-19 cells with exogenous interferon-b led to induction of ISG transcription, suggesting that ARPE-19 cells fail to sense or transduce early pathogen associated molecular patterns. Consistent with this hypothesis, ARPE-19 cells were found to express low steady-levels of the intracellular pattern recognition receptors CGAS and IFI16, as well as the Stimulator of Interferon Genes (STING). Consequently, interferon regulatory factors, such as IRF3, fail to acquire activating phosphorylation post-translational modifications, limiting their ability to transactivate interferon-b transcription. These data suggest that RPE cells generally fail to initiate interferon signaling in response to HCMV and/or other viruses, perhaps due to the eye being an immune privileged anatomical site; or that the ARPE-19 cell line may have lost the ability to respond to intracellular stresses during the spontaneous immortalization process. Our studies continue to highlight important characteristics of the ARPE-19 cell line that should be considered when using this cell line for biochemical or virological studies.

## Results

### 1.1 Comparative Proteomics Analysis of Infected MRC-5 and ARPE-19 Cells Identifies A Differential Intrinsic Immune Response to Human Cytomegalovirus

To identify cell-type specific responses to HCMV infection across fibroblasts and epithelial cells we conducted a whole-cell proteomics study. For this analysis, we used HCMV strain TB40/E-BAC4 and infected target cells at a multiplicity of five infectious units per cell, with the goal of infecting the majority of cells. Virus preparations were grown in ARPE-19 cells, which supports efficient entry into ARPE-19 and other pentamer-dependent cell lines^15,16,21,40–42^. Proteins were extracted from six samples (MRC5-mock, MRC-5-infected-replicate1, MRC5-infected-replicate2, ARPE-19-mock, ARPE-19-infected-replicate1, ARPE-19-infected-replicate2), at three time points (24, 72, and 120 hours post infection); and tryptic peptides were prepared for tandem mass-spectrometry analysis as described in “Materials and Methods.” Peptides from individual samples were labeled with isobaric tags (“tandem mass tags; TMTs), allowing for relative quantification of acquired spectra using reporter ions embedded in each MS2 fragmentation spectrum (Figure 1A). Data was analyzed using Isobar software^43^, which empirically models the noise between biological replicates to accurately assign significance values to differentially expressed proteins (Supplemental Figure 1).This type of modeling is important for accurate interpretation of quantitative mass-spectrometry measurements since reporter ions derived from isobaric tags exhibit a heteroscedastic signal-to-noise relationship^43^.

**Figure 1.**
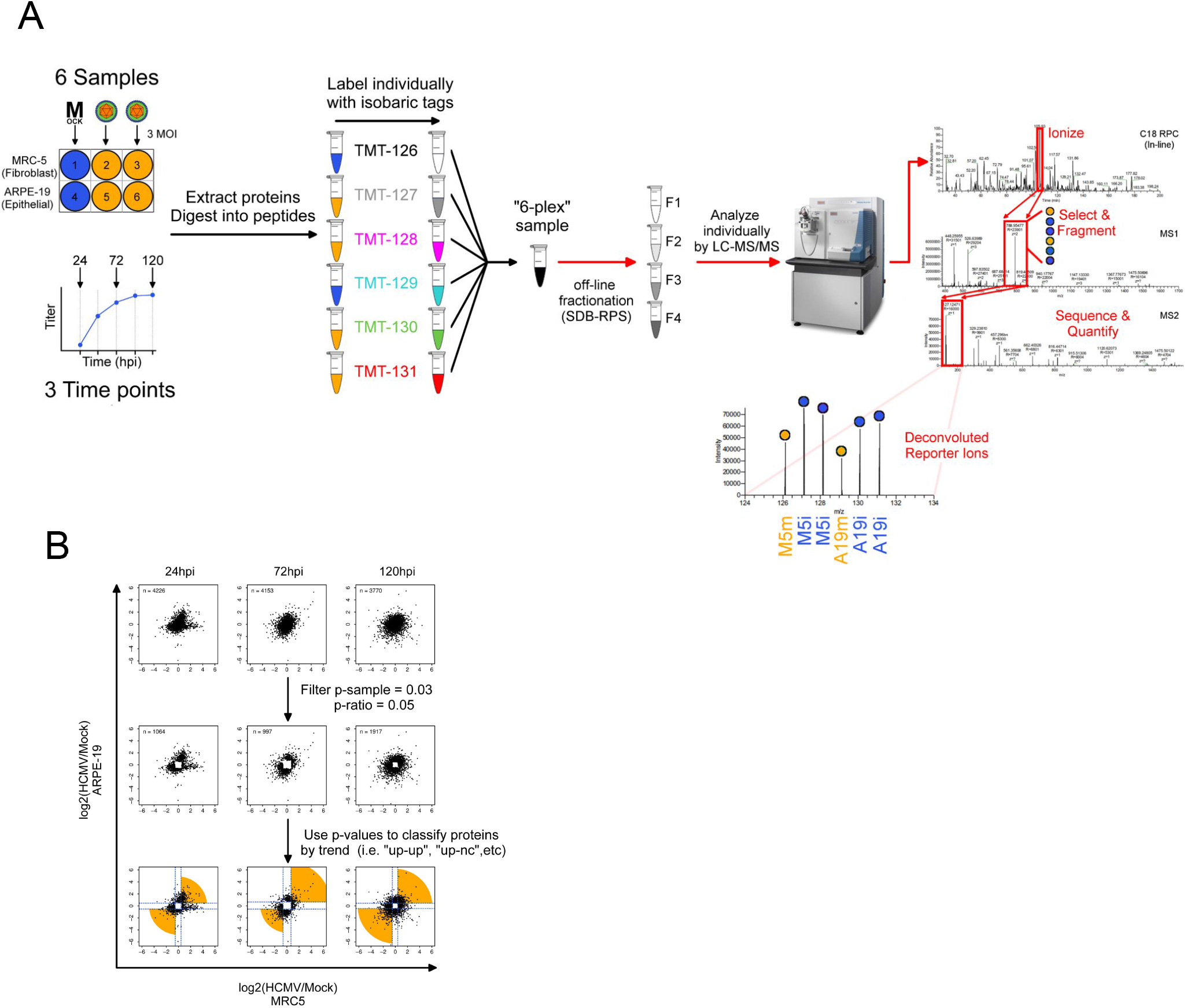
Whole-cell Mass-Spectrometry Workflow for Comparative Analysis of HCMV-infected MRC5 and ARPE-19 cells. (A) MRC5 fibroblasts and ARPE-19 cells were mock infected or infected with TB40/E-BAC4 at 3 MOI. Whole-cell lysates were collected at 24, 72, and 120 hours post infection (hpi) and prepared for tandem mass-spectrometry analysis using isobaric tags for relative quantification (see methods). (B) Scatter plots depicting infected/mock ratios of cellular proteins identified by MS/MS analysis across MRC5 and ARPE-19 cells. Proteins differentially regulated in both cell types, or differentially regulated in one cell type but not the other, were identified using a two step strategy. First, summary statistics (p-values) were used to filter insignificantly regulated proteins. Second, remaining proteins were grouped by trend (e.g. “up-up”, “up-nc”, etc.) for further analysis. *nc = no-change*.

Isobar assigns two p-values to each protein: a “ratio” p-value inversely correlating to the pooled spectrum quality of an observed protein and a “sample” p-value indicating the inversely correlating to the confidence of differential expression of a given protein across two samples; as well fold-change ratios for each observed protein (Supplemental Figure 1).

We initially set out to identify cellular proteins that differentially responded to HCMV infection across MRC-5 fibroblasts and ARPE-19 retinal pigment epithelial cells. We hypothesized that, given the biological differences between fibroblasts and epithelial cells, different pathways might be activated or inhibited during infection of each cell type. Therefore, to experimentally address this hypothesis we analyzed proteomic changes across the two cell types, over a five days infection time course, anticipating the identification of differential single-factor or pathway-level changes indicative of differential cellular responses to infection. Our general analytical strategy involved: (1) assigning infected/uninfected ratios to each identified protein in each cell type; (2) assigning identified proteins to eight classes based on their fold-change trends and statistics (e.g. “A19-up/M5-up”, “A19-no-change/M5-up”, “A19-down/M5-down”, etc.); and (3) performing enrichment analysis on each class to identify groups of proteins suggestive of pathway-level activation, or inhibition, in each class (Figure 1B).

Using this procedure, we identified a group of related proteins that were strongly upregulated in MRC-5 cells at 24 hpi, but did not change significantly in ARPE-19 cells (Figure 2A); a putative cell type specific response. For this analysis, we considered a protein significantly regulated if its p-ratio was less-than-or-equal to 2-fold and its p-sample was less-than-or-equal to 0.03. The proteins in the MRC5-up, ARPE-19-no-change (NC) group constituted the most prominent cell-type specific response we could identify across MRC-5 and ARPE-19 cells using this approach.

**Figure 2.**
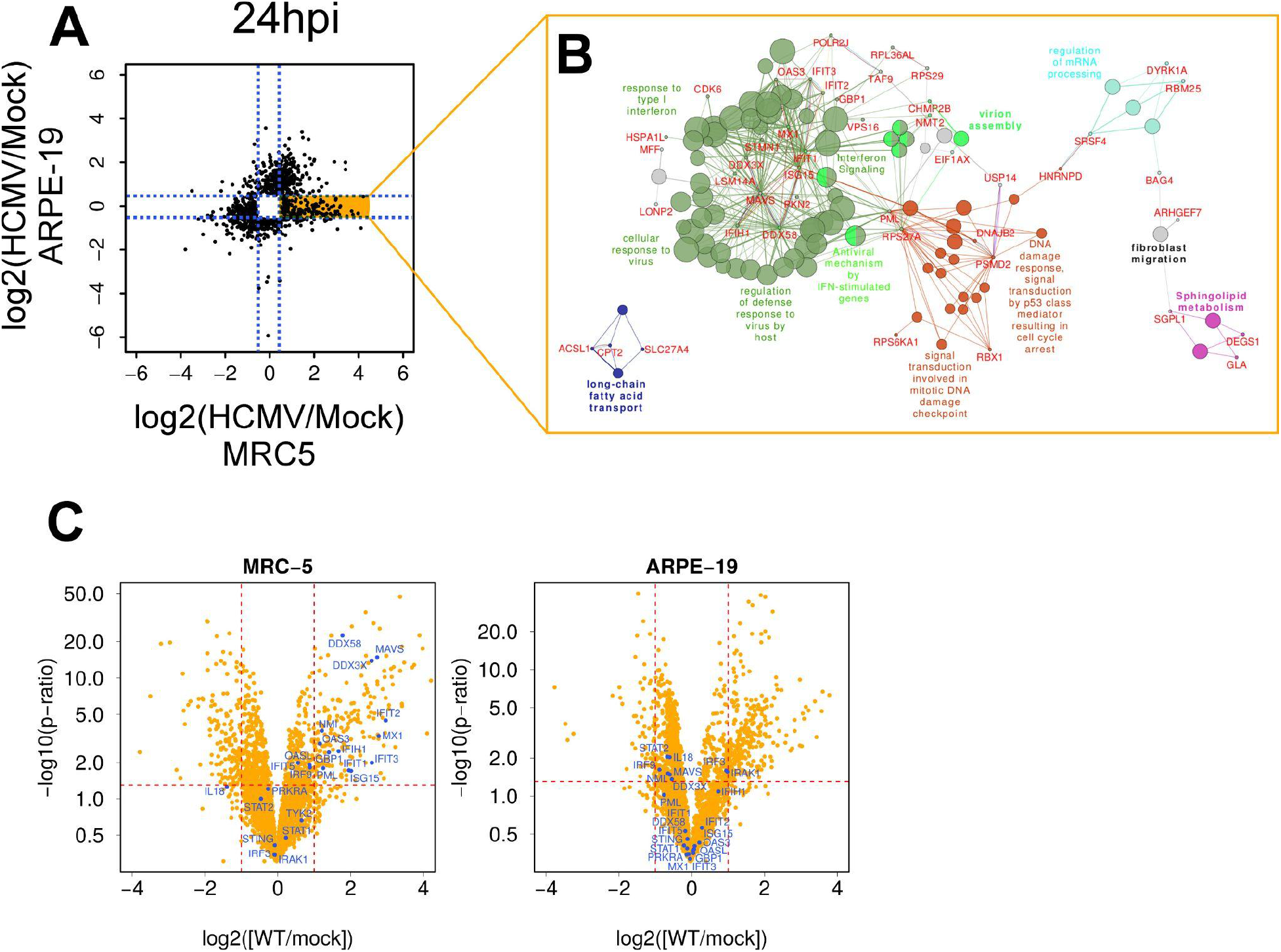
Interferon Stimulated Genes (ISGs) are Activated in MRC5 Cells, but Not ARPE-19 Cells, After Infection with HCMV. (A) Isobar p-values were used to identify proteins upregulated by infection in MRC5 cells, but unchanged by infection in ARPE-19 cells. (B) Enrichment analysis was performed using the Cytoscape plugin Cluego. Displayed is a combined ontology network and protein interaction network where large circular nodes represent enriched ontology terms, small circular nodes represent proteins, and edges represent interconnectedness. A large community of enriched interferon-stimulated ontology terms and proteins (green circles) suggest type-I interferon signaling is initiated in MRC5 cells, but not ARPE-19 cells, after infection with HCMV.

Enrichment analysis using the Cytoscape^44^ plugin Cluego^46^ identified a significant enrichment in a protein-set associated with the response to Type I interferon (Figure 2B). Mapping gene/protein identifiers from the MSigDB^45^ geneset “Hallmark Type-I Interferon Response” back to the proteomics dataset at 24 hpi, in both MRC-5 cells and ARPE-19 cells revealed that ISGs such as IFIT1, IFIT2, IFIT3, MX1, OAS3, and ISG15 were strongly and coordinately upregulated at the protein level in MRC-5 cells, but were not significantly changed in ARPE-19 cells upon infection with HCMV (Figure 2C). These changes were validated by analyzing a previously acquired RNA-sequencing data set^40^ of cellular mRNA levels across HCMV-infected MRC5 and ARPE-19 cells (Figure 3A) and independently performing *q*RT-PCR for a subset of ISGs (Figure 3B). mRNA-level changes of ISGs were linearly correlated (R^2 0.54) with protein level changes in MRC-5 cells; while no evidence for ISG induction in ARPE-19 cells was observed in any experiment.

**Figure 3.**
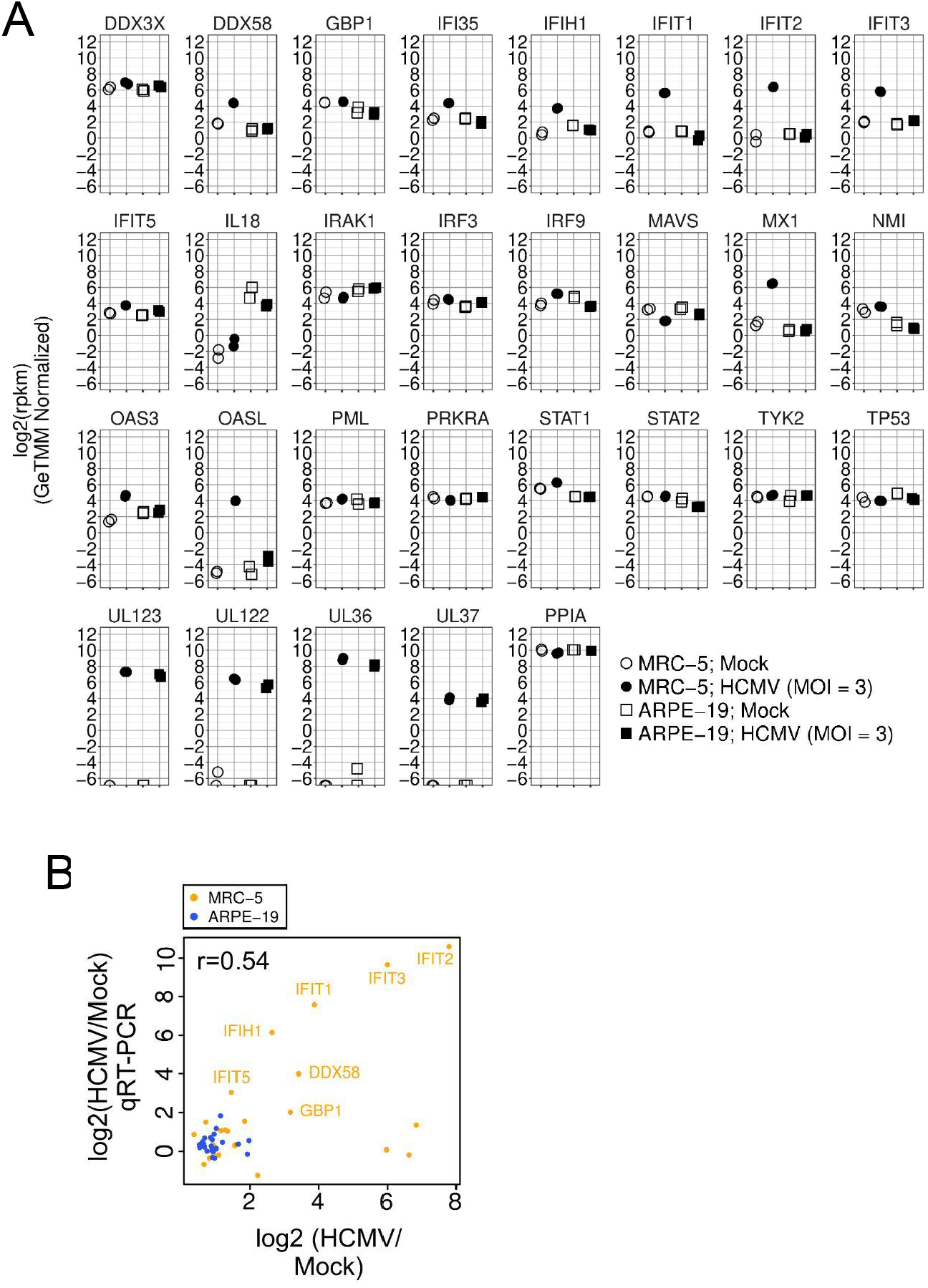
Validation of MS Data Using RNA-sequencing and *q*RT-PCR Analysis. RNA-sequencing data from our previous study (Oberstein and Shenk, 2017, PNAS) was re-analyzed. Absolute mRNA levels for a set of interferon stimulated genes are displayed. mRNAs for *DDX58, IFI35, IFIT1, IFIT2, IFIT3, MX1, OAS3*, and *OASL* show upregulation in HCMV-infected MRC5 cells, but not ARPE-19 cells; similar to our independent MS observations. *PPIA* and *RPLP0* are displayed as normalization controls. The HCMV immediate-early transcripts *UL123, UL122, UL36, and UL37* mark HCMV-infected samples. Data represent the absolute expression level in reads-per-kilobase-per-million (rpkm) in each cell-type, after cross-sample normalization using the GeTMM procedure (see “Material and Methods”). (B) Correlation of ISG mRNA (*q*RT-PCR) and protein (MS) levels in HCMV-infected MRC5 (yellow points) and ARPE-19 (blue points) cells. ISG mRNAs and proteins showed a moderately good linear correlation in MRC5 cells (spearman’s coefficient of 0.54); while neither ISG mRNAs or proteins were significantly changed in HCMV-infected ARPE-19 cells.

### 1.2 Effect of UV-inactivation of HCMV on the Type-I Interferon Response in MRC-5 and ARPE-19 Cells

Like many viruses, HCMV expresses early gene products that antagonize type-I interferon signaling. To determine whether the lack of interferon response in ARPE-19 cells was due to cell-type specific expression of a viral gene product or a general defect in host cell initiation of interferon signaling we performed infections with UV-inactivated HCMV across ARPE-19 cells and control MRC-5 cells. UV-inactivation inhibits *de novo* gene expression from the incoming HCMV genome and interferon-competent cells typically show increased expression of interferon-b and ISGs in the presence of UV-inactivated HCMV, since interferon antagonistic viral gene products fail to be expressed^47–50^. As expected, equal innocula of untreated and UV-treated HCMV both initiated robust expression *IFNB1* and ISG mRNA in MRC-5 fibroblasts (Figure 4), with *IFNB1* increasing approximately 10-fold and ISGs increasing between 2 - 8-fold after infection with UV-inactivated HCMV (Figure 4). However, neither untreated or UV-treated HCMV induced transcription of *INFB1* or downstream ISG mRNAs in ARPE-19 cells (Figure 4). Therefore, we conclude that the lack of interferon production in ARPE-19 cells is independent of *de novo* HCMV gene expression.

**Figure 4.**
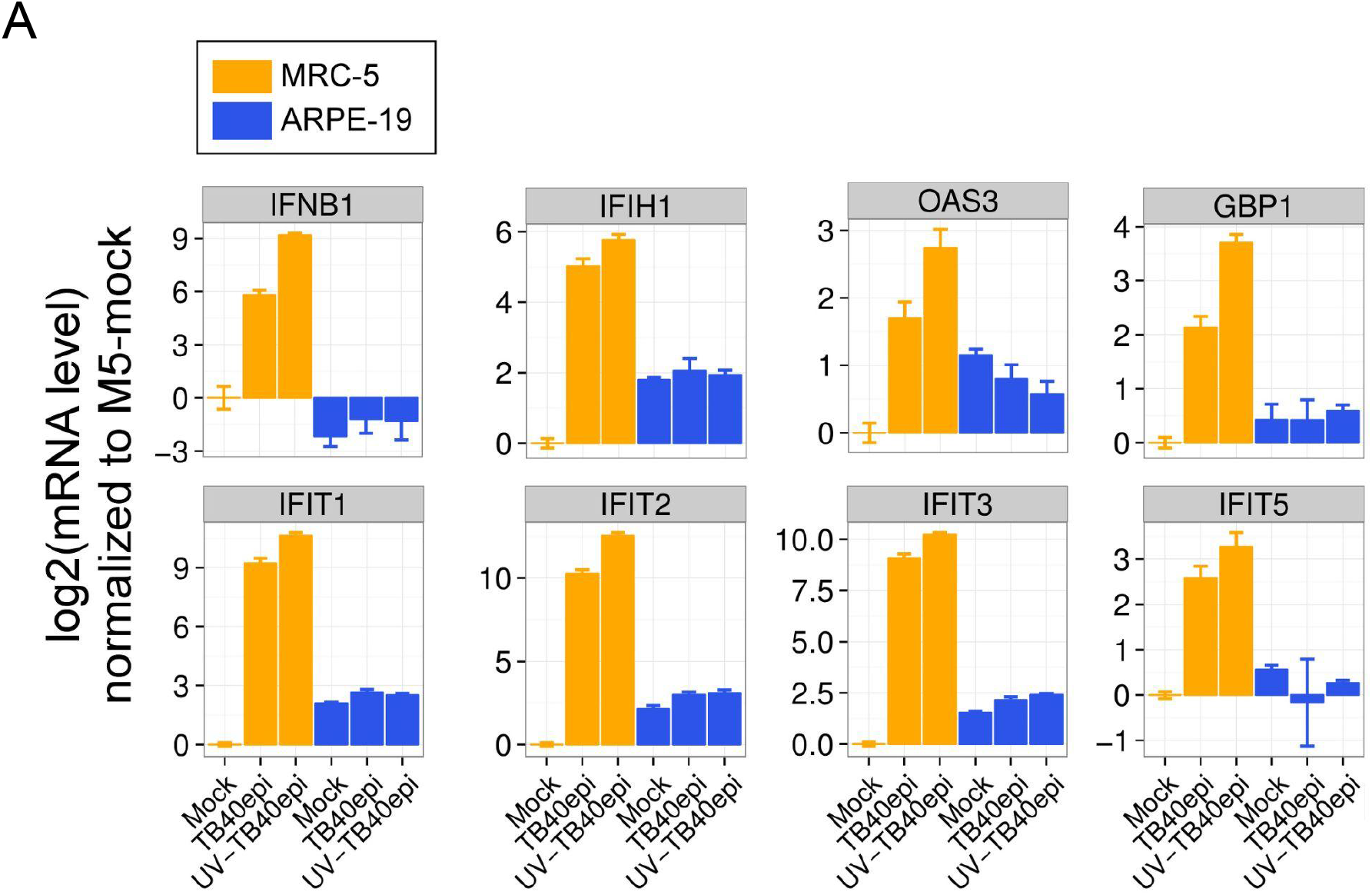
The Lack of Type-I Interferon Signaling in ARPE-19 cells is Independent of de novo HCMV Gene Expression. Relative mRNA levels of interferon-beta (*IFNB1*) and select ISGs across MRC5 and ARPE-19 cells infected with untreated (TB40epi) and UV-inactivated (UV-TB40epi) HCMV. an identical stock of TB40/E-BAC4 produced in ARPE-19 cells (TB40epi) was UV-inactivated and used to infect side-by-side cultures of MRC5 and ARPE-19 cells at 3 MOI. RNA was extracted from mock and infected samples at 24 hpi and analyzed by qRT-PCR. Data represent the mean +/−SD from three biological replicates assayed in technical triplicate.

### 1.3 Effect of HCMV Growth Host and Recombinant Interferon on the Type-I Interferon Response in MRC-5 and ARPE-19 Cells

To further characterize the mechanism underlying the differential interferon response across MRC-5 and ARPE-19 cells, we assessed the impact of HCMV growth host and the capacity for both cell types to respond to interferon treatment. For this, we monitored *IFNB1* and ISG transcription using *q*RT-PCR after treatment with HCMV produced in both ARPE-19 and MRC5 cells, as well as after treatment with recombinant Interferon-b. A *IFNB1* and ISGs were efficiently induced in MRC-5 cells, but not ARPE-19 cells irrespective of whether HCMV was produced in fibroblasts or epithelial cells (Figure 5). In addition, treatment with recombinant human Interferon-b efficiently induced ISG transcription to a similar magnitude in both cell types. Thus, ARPE-19 cells are capable of initiating ISG transcription in the presence of Type-I interferons, but do not induce ISG transcription in response to HCMV infection. These results suggest that the failure to initiate Type-I interferon signaling in ARPE-19 cells is likely due to an upstream lesion in HCMV sensing, signal transduction, or activation of *IFNB1* transcription; rather an inability to respond to secreted interferon or downstream JAK-STAT signaling.

**Figure 5.**
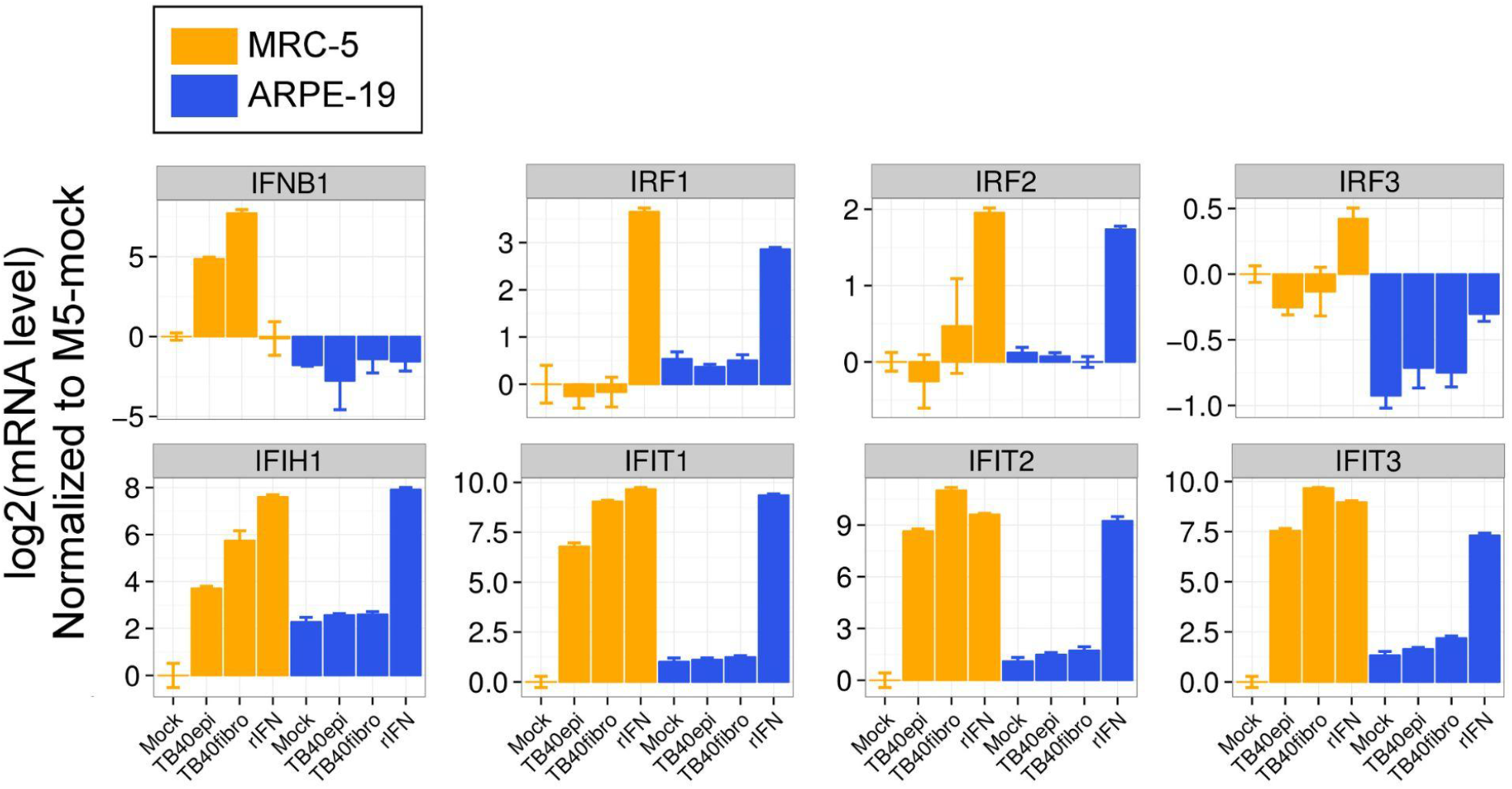
ARPE-19 Cells are Competent to Respond to Exogenous Type-I Interferon, but Fail to Initiate Interferon Signaling, Irrespective of HCMV Growth Host. Relative mRNA levels of interferon-beta (*IFNB1*) and select ISGs across MRC5 and ARPE-19 cells infected with TB40/E-BAC4 produced in ARPE-19 cells (TB40epi) or MRC5 fibroblasts (TB40fibro); or after treatment with recombinant human Interferon-b in the absence of infection (rIFN). Infections were performed at 3 MOI. RNA was extracted from all treatments at 24 hpi or 24 post-treatment (rIFN), and analyzed by *q*RT-PCR. Data represent the mean +/−SD from three biological replicates assayed in technical triplicate.

### 1.4 ARPE-19 Cells Express Low Levels of IFI16, CGAS, and STING and Fail to Activate IRF-3 Phosphorylation Upon Infection with HCMV

The intrinsic immune response to HCMV in productive fibroblasts involves recognition by pattern recognition receptors and signal transduction resulting in activation of inducible transcription factors responsible for activating type-I interferons. Canonically, this involves phosphorylation of interferon regulatory factors (IRFs), relocalization to the nucleus or allosteric activation of DNA-binding activity, and direct or indirect recruitment of RNA-polymerase II to the interferon gene promoters. Intracellular DNA-sensing pathways responsible for sensing foreign, or mislocalized, nucleic acids^51–61^, as well as pattern recognition receptors on cell surface or endosomal membranes (i.e. Toll-like receptors (TLRs))^62–70^, initiate the intrinsic immune response to cytomegaloviruses. Although ARPE-19 cells are not immortalized, selective trypsinization and serial passaging was used to establish the cell line^20^, which may have selected for alterations in proteins or pathways important for initiating the intrinsic immune response to foreign nucleic acids. Given our observations with HCMV and recombinant IFNb, we hypothesized that the cell-culture conditions used to derive the ARPE-19 cell line may have selected for a lesion in intrinsic DNA-sensing, and/or early interferon signal transduction, upstream of *IFNB* transcription. Alternatively, retinal pigment epithelial cells may have an naturally low intrinsic response to viruses; perhaps due to the eye being an “immune privileged” anatomical site^71^. Therefore, we hypothesized that ARPE-19 cells might exhibit low expression of one or more critical DNA-sensors or effectors. Therefore, we assessed the expression levels of cytosolic DNA sensors CGAS^53,57^, IFI16^60^, and ZBP1^48^, each of which have been shown to sense HCMV during infection; as well as the critical signal transduction protein STING^56,58^, across *uninfected* ARPE-19 and MRC-5 cells. mRNA levels of *CGAS* were approximately 4-fold lower in uinfected ARPE-19 cells than MRC5 cells; while mRNA levels of *IFI16* and *STING1* were approximately 16-fold lower in ARPE-19 cells than MRC5 cells (Figure 6A). mRNA levels of *ZBP1* showed no significant difference between the cell lines (Figure 6A). These results were confirmed by re-analyzing RNA-sequencing data from our previous transcriptomics study of HCMV-infected MRC-5 and ARPE-19 cells (Figure 6B; ^40^). Consistent with these results, IFI16 protein was nearly undetectable in ARPE-19 cells by immunoblot, while both CGAS and STING steady-state protein levels were reduced approximately 3-fold and 6-fold, respectively, compared to MRC5 cells (Figure 6C). ZBP1 protein was expressed at similar levels in both ARPE-19 and MRC-5 cells (Figure 6C); again consistent with the mRNA expression data. Not surprisingly, phosphorylation of IRF3, a critical trans-activator of the *IFNB* promoter, was reduced approximately 4-fold in HCMV-infected ARPE-19 cells compared to infected MRC-5 cells. These data support a model by which low endogenous levels of cytosolic DNA-sensors and/or the signal transducer STING disable early sensing of immunogenic features of incoming HCMV virions or early interferon signal transduction (e.g. STING activation) responsible for IRF3-phosphorylation (Figure 7). Consequently, *IFNB* transcription, autocrine and paracrine activation of type-I interferon receptors, JAK-STAT signaling, and ISG induction fails to occur in response to HCMV infection in ARPE-19 cells (Figure 7).

**Figure 6.**
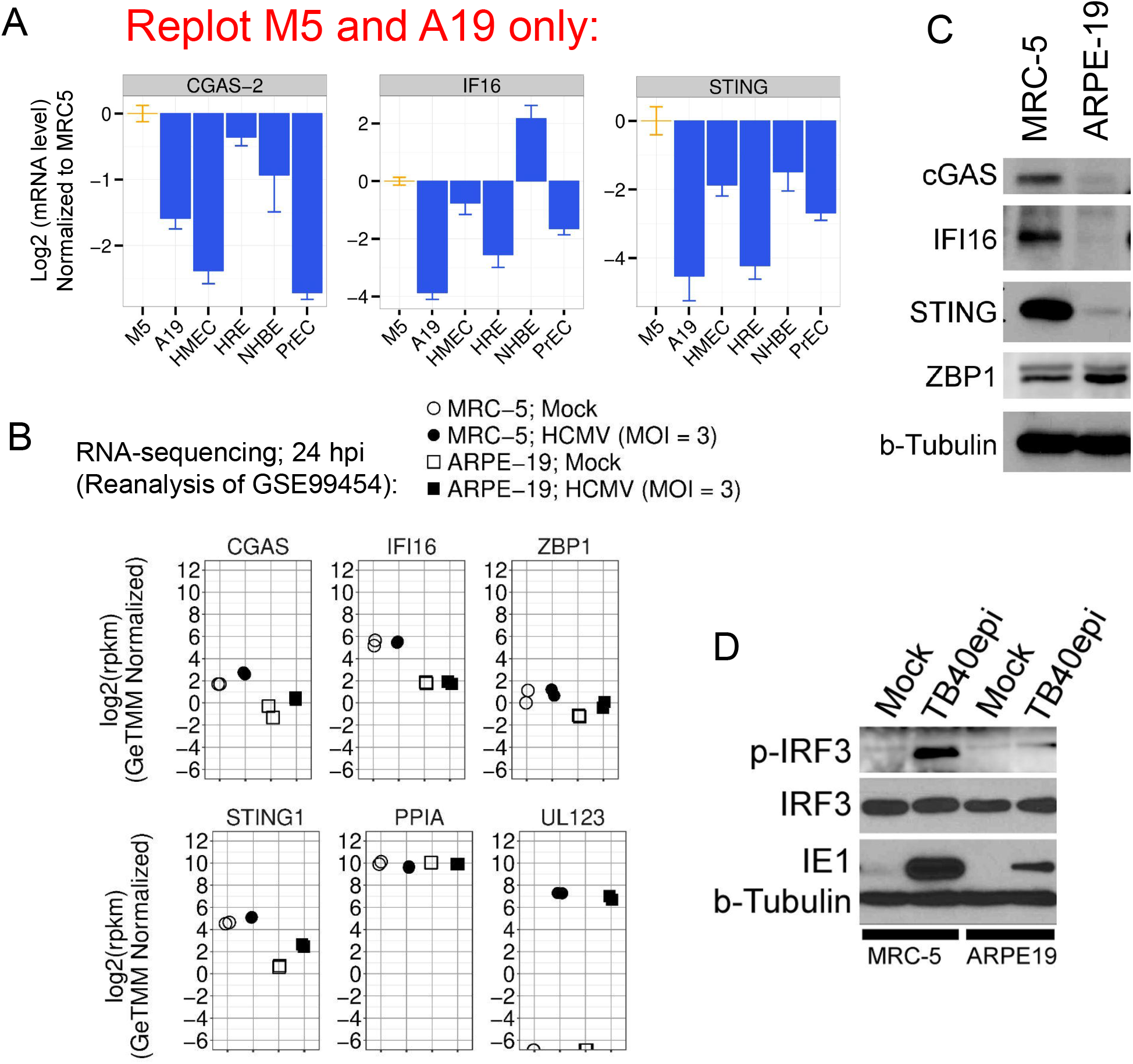
ARPE-19 Cells Express Low Levels of IFI16, CGAS, and STING and Fail to Activate IRF-3 Phosphorylation Upon Infection with HCMV. (A) Relative mRNA levels of *CGAS, IFI16, and STING* across MRC5 and ARPE-19 cells, measured by qRT-PCR. Data represent the mean +/−SD from three biological replicates assayed in technical triplicate. (B) Re-analysis of public gene expression data from GSE99454 (Oberstein and Shenk, PNAS, 2017). Relative mRNA levels of *CGAS, IFI16, STING*, and *ZBP1* are displayed for MRC5 and ARPE-19 cells, in the presence and absence of HCMV infection. The viral immediate-early mRNA, *UL123*, and cellular mRNA *PPIA* are shown as infection and normalization controls, respectively. (C) Immunoblot (western) analysis showing steady-state protein levels of IFI16, CGAS, STING, and ZBP1 across MRC5 and ARPE-19 cells. b-Tubulin is shown as a loading control. (D) Immunoblot showing relative phosphorylation levels of IRF3 after HCMV infection across MRC5 and ARPE-19 cells. Total IRF3, HCMV IE1, and b-Tubulin are displayed as controls.

**Figure 7.**
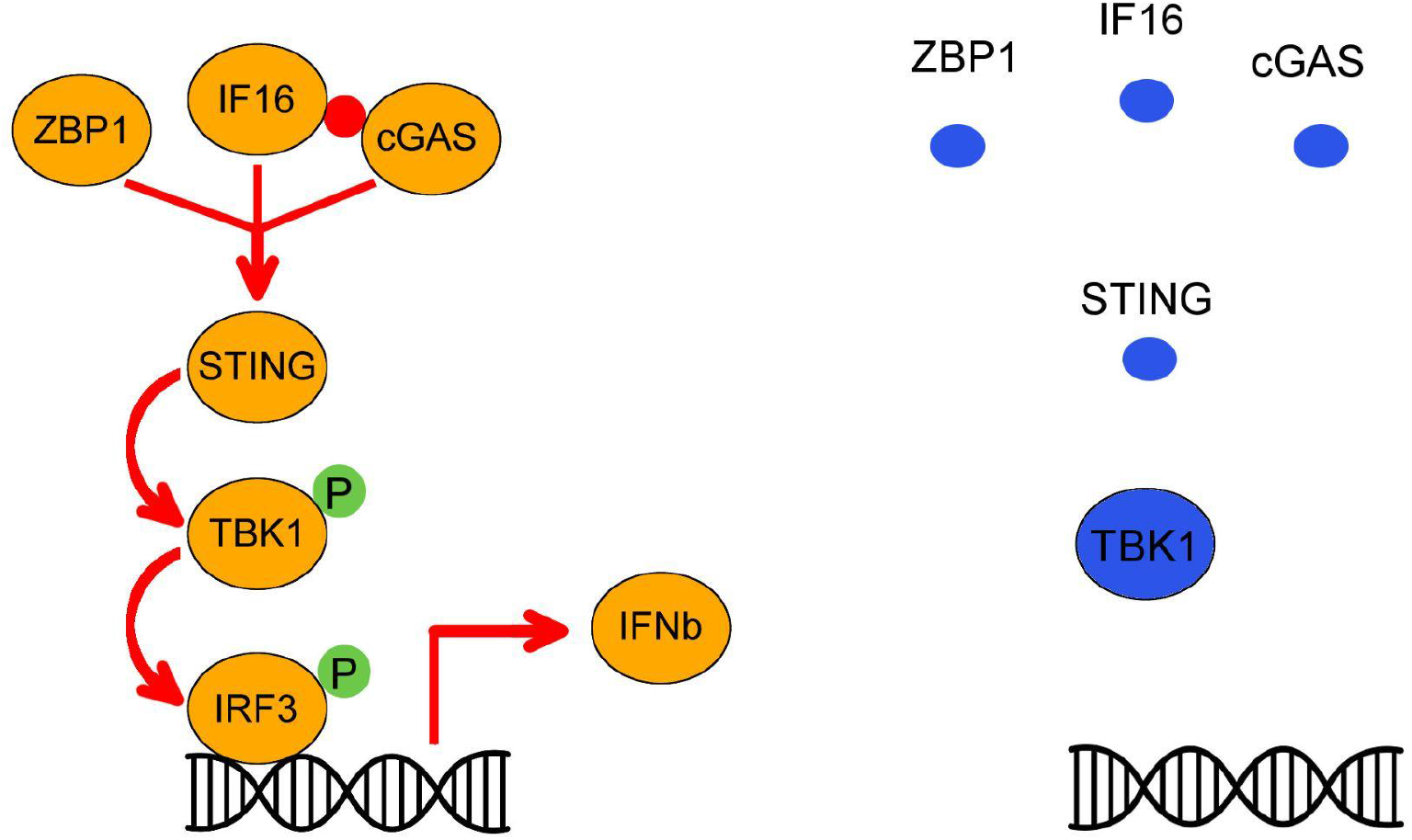
Cytosolic Sensor Model for Explaining Lack of Type-I Interferon Signaling in ARPE-19 Cells. Cytosolic sensing of HCMV occurs normally in MRC5 cells (left; orange circles), leading to efficient activation of *IFNB1* transcription and downstream signaling leading to a canonical type-I interferon response. Low endogenous levels of cytosolic DNA-sensors and the signal transducer STING in ARPE-19 cells (right; blue circles) disable sensing of HCMV virions and STING activation. Consequently, IRF3 phosphorylation and *IFNB1* transcription fail to occur, blunting downstream interferon signaling including autocrine and paracrine activation of type-I interferon receptors, JAK-STAT signaling, and ISG activation. Size of proteins/circles is proportional to their relative amounts.

## Discussion

In this study we set out to determine cell-type specific responses to HCMV across fibroblasts and ARPE-19 epithelial cells. HCMV infects a wide array of cell types in the body. An important outstanding question in CMV biology is whether different cell types initiate different responses to HCMV infection, especially since the virus is thought to establish both quiescent (latent) and productive biosynthetic states in different cell types. Thus far, the proteomics data presented in this study show very few differential responses to HCMV-infection across fibroblasts and ARPE-19 cells, at either early or late times after infection. One exception was the type-I interferon response, which we thoroughly validated using a variety of methods.

ARPE-19 cells are a commonly used epithelial cell culture model used for studying retinal epithelial biochemistry and viral infection. Although ARPE-19 cells recapitulate some aspects of RPE biology^20,27^, they require long-periods of high density “differentiation” to attain structural and biochemical characteristics of primary RPE cells ^20,27^.

HCMV naturally infects retinal epithelial cells and primary infection or reactivation from latency can present in the eye, causing retinitis; especially in immunocompromised carriers such as HIV/AIDS patients. ARPE-19 cells have been used extensively to model HCMV infection *in-vitro*, as they are generally thought to provide a physiologically relevant, reliable, and renewable source of RPE-like cells for infection studies.

Our work has uncovered several aspects of the ARPE-19 cell line that should be considered when using ARPE-19 cells for *in-vitro* studies with HCMV and/or other human viruses. First, low-confluency cultures of ARPE-19 cells assume a mesenchymal cell state phenotype, which could affect one or more aspects of the HCMV infectious cycle^21^. A comparison of HCMV progeny production in “mesenchymal” ARPE-19 cells and other strongly epithelial cells lines suggests that the epithelial-mesenchymal cell state of ARPE-19 cells may influence HCMV biosynthesis and progeny production^21^. Second, we now find that subconfluent ARPE-19 cells lack a type-I interferon response to HCMV infection, possibly due to low expression levels of CGAS, IFI16, and STING. Whether or not this lack of capacity to initiate an interferon response to HCMV is due to a lesion created during the establishment of the ARPE-19 cell line or whether RPE cells generally lack the ability to initiate interferon signaling, remains unclear. However, the inability to initiate interferon signaling in response to HCMV may limit the use of ARPE-19 in some studies, particularly those looking at the cellular response to infection.

Both primary RPE and ARPE-19 cells are capable of inducing *IFNB* in response to Poly I:C^72^, transfected RNA^73,74^, or poly-dA-dT^74^, a synthetic form of B-DNA^75–77^. However, previous work with ARPE-19 cells showed a severely limited interferon response due to transfected dsDNA^74^. The ARPE-19 response to RNA appears dependent on the RNA-sensor RIG-I/DDX58^73,74^. Surprisingly, a clear comparison of the type-I interferon response between primary RPE and ARPE-19 cells infected with HCMV seems to be absent from the literature. Future studies comparing primary RPE and ARPE-19, appropriately differentiated to maintain epithelial characteristics, are needed to fully characterize ARPE-19 cells. Until then, we continue to urge caution when interpreting studies using ARPE-19 cells as a general epithelial model for use with HCMV or other human pathogens.

## Material and Methods

### Cells, Viruses, and Plasmids

MRC-5 human embryonic lung fibroblasts and ARPE-19 adult retinal pigment epithelial cells were obtained from the American Type Culture Collection and grown in DMEM supplemented with 10% fetal bovine serum, 1 mM sodium pyruvate, 2 mM glutamax (Gibco), 10 mM Hepes pH 7.4, 0.1 mM MEM Non-Essential Amino Acids (Gibco), 100 units/mL Penicillin G, and 100 μg/mL Streptomycin Sulfate. HCMV strain TB40/E-BAC4^78^ was reconstituted by electroporation into ARPE-19 cells, expanded on 15 cm dishes of ARPE-19 cells, and concentrated by ultracentrifugation over a 20% sorbitol cushion. Virion-containing pellets were resuspended in PBS containing 7% sucrose and 1% BSA, aliquoted, and stored at −80° C until use. Virus preparations were titered on both MRC-5 and ARPE-19 cells using an HCMV infectious unit assay, as previously described^21^.

### Quantitative reverse transcription polymerase chain reaction (qRT-PCR)

RNA was isolated using Trizol reagent (Ambion) followed by purification using Qiagen Rneasy columns. Contaminating DNA was removed using Turbo Dnase (Thermo Fisher Scientific) and 1 ug total RNA was reverse-transcribed in a 20 ul reaction volume using a Superscript VILO cDNA synthesis kit (Invitrogen). *q*RT-PCR reactions were performed in 384-well plates in a 10 ul reaction volume by mixing 0.7 - 1.7 ul of 10-fold diluted cDNA from each sample with 0 – 1.0 ul H20, 5 ul 2x Power SYBR Green Master Mix (Applied biosystems) and 3.3 ul 1.2 uM PCR primers ([primer]final = 0.4 uM). qRT-PCR reactions were performed using Power SYBR Green Master Mix (Applied biosystems) and data were collected on a Viia7 digital PCR machine (Life Technologies). Data were analyzed using the ΔΔCT method^79^ with cyclophilin A (PPIA) as the reference gene. All primers were designed using either QuantPrime^80^ or GETPRime^81^. *q*RT-PCR primer sequences are listed in Supplementary Table S2.

### Western Blotting

Cell monolayers were collected in RIPA buffer (50 mM Hepes pH7.4, 150 mM NaCl, 1% NP-40, 0.5% sodium deoxycholate, 0.1% SDS, 5 μg/mL aprotinin, 10 μg/mL leupeptin, 1 mM PMSF). Samples were then sonicated and debris was cleared by centrifugation. Protein concentration was determined using the BCA assay (Pierce) and equal amounts of total protein were separated by SDS-PAGE. Proteins were transferred to PVDF membranes and blocked with 5% BSA in HBST (50 mM Hepes pH 7.4, 150 mM NaCl, 0.05% tween-20). Primary antibodies (Supplementary Table S3) were diluted in 1% BSA and rocked at 4 °C overnight. Primary antibodies were detected using HRP-conjugated secondary antibodies and ECL Prime (Cytiva). Antibodies and dilutions are listed in Supplementary Table S3.

### Experimental Infections

One day prior to infection, cells were seeded into complete medium at 3×10^5^ per well (MRC-5) or 2×10^5^ per well (ARPE-19) in 6-well dishes. For mass-spectrometry analysis, two independent infections were performed (biological replicates) with a single common “mock” infected well, for each cell type, at each time point (see Three independent infections were performed per cell type for each time point. Both cell types had just reached confluency at the time of infection. Unless otherwise indicated, cells were infected at 3 infectious units per cell (IU/cell) by adding an appropriate HCMV inoculum to the cell culture supernatant for 2 h in a standard tissue culture incubator at 37°C and 5 % CO_2_. After the infection period, cells were washed one time with warm PBS and before returning the infected cells to the tissue culture incubator in complete medium.

### Preparation of Peptides for Tandem Mass-Spectrometry Analysis

At each time point, tissue culture plates were placed on ice, washed one time with ice-cold PBS, and scraped into MS Lysis buffer (0.1 M Ammonium Bicarbonate, 2.5 % Sodium Deoxycholate, 5 mM Sodium Orthovanadate, 50 mM Sodium Fluoride). Lysates were then flash frozen in liquid nitrogen and thawed one time simultaneously once all time points had been collected. Side-by-side infections were collected in Trizol reagent (Ambion) for qRT-PCR analysis.

Lysates were thawed, sonicated for 10 pulses using a tip sonicator, and quantified using a BCA assay (Pierce). 100 ug of total protein for each sample was then adjusted to a total volume of 300 ul with MS Lysis buffer, reduced with 5 mM Tris(2-carboxyethyl)phosphine hydrochloride (TCEP; Pierce) for 20 min at 55° C, and alkylated with 10 mM Chloroacetamide (Sigma) for 30 min at 37° C. Proteins were then digested into peptides by the addition of 1:1000 (g/g) Lys-C Protease (Promega) for 4 h at 37°C, followed by the addition of 1:50 (g/g) sequencing grade Trypsin (Promega) for 16h at 37° C. Proteases were inactivated by the addition of 1 mM PMSF and 10 ul 1x complete protease inhibitor cocktail (Roche).

Peptides were phase extracted to remove detergent (deoxycholate) by adding 1.0 ml pure Ethyl Acetate (Sigma), followed by 15 ul 20 % Trifluoroacetic Acid (TFA), rapid mixing using a vortexer, and centrifugation. The detergent containing organic upper phase was removed and traces of Ethyl acetate were removed using a speedvac. Finally, peptides from each sample were purified using a 1 ml Sep-pak C18 column (Waters), concentrated to approximately 50 ul using a speedvac and quantified using a BCA assay (Pierce).

### Tandem-mass tag (TMT) Labeling and Fractionation

TMTsixplex Isobaric Label Reagent kit (Pierce) was used to multiplex samples at each time point (24, 72, and 120 hpi). Each 6-plex sample consisted of: MRC5-mock (TMT-126), MRC5-HCMV1 (TMT-127), MRC5-HCMV1 (TMT-128), ARPE19-mock (TMT-129), ARPE19-HCMVrep1 (TMT-130), ARPE19-HCMVrep2 (TMT-131). For TMT-labeling, 10 ug of peptides from each sample were adjusted to pH 8.5 by addition of 1/10 V 1.0 M Triethylammonium bicarbonate (Pierce) and labeled with the appropriate TMT-reagent by adding 0.2 mg TMT-reagent solubilized in 100 % Ethanol for 1h at room temperature. Reactions were quenched by the addition of 5 % hydroxlamine (Sigma). 6-plex samples were then pooled and partitioned into four fractions using Empore SDB-RBS (3M) stage tips^82^. Peptides were eluted using the following buffers:

1. 0.15 M ammonium formate, 0.5% FA, 40% ACN
2. 0.20 M ammonium formate, 0.5% FA, 60% ACN
3. 0.20 M ammonium acetate, 0.5% AA, 60% ACN
4. 5% ammonium hydroxide, 80% ACN

Finally, each fraction, from each sample, was vacuum-dried vacuum-dried to near dryness, and adjusted to 20 ul with 1% formic acid + 5% ACN.

### Mass-spectrometry data collection

Peptides were analyzed by nanoLC-MS on an LTQ-Orbitrap Velos mass-spectrometer (Thermo Fisher Scientific). Peptides were separated by reverse-phase C18 chromatography using a 90 min gradient on an in-line Dionex Ultimate nRSLC and ionized by electro-spray ionization (ESI) using a voltage of 1.8 kV and ion transfer tube temperature of 300 °C. For each run, 4 μL digest was loaded, and each sample was analyzed in technical triplicate. The instrument was set to acquire spectra in data dependent acquisition mode (DDA) with the top 10 most intense ions being selected for HCD fragmentation. Each acquisition cycle comprised a single full-scan mass spectrum (m/z = 385 – 1700) in the orbitrap (r = 30,000 at m/z = 400) followed by HCD MS/MS (fixed first m/z = 100) of the top 10 most intense ions (minimum signal = 2000). Dynamic exclusion was enabled (90 secs) and FT preview scan was disabled.

### Mass-spectrometry data analysis

Raw MS data (.raw files) was converted to mzXML format using msconvert^83^. Tools from the Trans-proteomic pipeline (TPP) software suite Version 4.7.1^84,85^ were used for database searching, peptide false-discovery rate modeling, and extracting TMT reporter ion-intensities. In-house python scripts were used to automate most processing steps. Database searches were performed individually on each of the three technical runs, per time-point, using two search algorithms, X!Tandem^86,87^ and Comet^88^, yielding six pep.xml files (6 pep.xml x 3 time-points x 2 virus strains = 36 searches total). For both algorithms semi-tryptic peptides were allowed, the precursor ion tolerance was set at 5 ppm, and modifications were set to require cysteine alkylation (static: +57.021464), TMT modifications on the N-terminus and Lysine (static: +229.162932 Da), possible Methionine oxidation (variable: +15.9949), and possible deamidation of Asparagine (variable: +0.984016). Search databases contained concatenated protein sequences from the uniprot human reference proteome (canonical Swiss-Prot only as of January 2016), common FBS and molecular biology (Trypsin, GFP, etc.) contaminants, and all canonical HCMV proteins from strain TB40/E-BAC4^78^. Randomized decoy versions of all sequences were appended to the search databases for target-decoy modeling. X!Tandem results were converted to pep.xml^84^ using the tpp function tandem2xml, while comet directly outputs a .pep.xml. Initial identification probabilities were generated for each search algorithm separately by pooling all identified peptides from the triplicate technical replicates and modeling correctly vs incorrectly (“decoy”) assigned spectra using PeptideProphet^89^. Identification probabilities were refined by using InterProphet^90^ to merge and rescore the PeptideProphet results from the two search engines. Isotope corrected, reporter ion intensities were extracted from each spectrum using libra^91^. Spectra were filtered to a final protein level FDR of 1% using MAYU^92^. For all datasets this amounted to a peptide level FDR well below 1% (see complete dataset stats in Supp Table XX).

Isobar software^43^ was used to generate protein level summary statistics for treatment contrasts (i.e. fold-change ratios and significance measures). Isobar uses an empirical noise model between same-same “intra-class” ratios to accurately assign statistical significance to “inter-class” ratios (e.g. HCMV/mock ratios in each cell type, at each time point). Isobar’s data model provides two p-values: p-ratio, a measure of ratio quality, and p-sample, which is classical statistical significance determined by testing each “interclass” ratio ([Avg(HCMV/mock)]proteinX) against an experimentally derived null-distribution consisting of all pooled “intraclass” ratios (MRC5-HCMV1/MRC5-HCMV2 + ARPE19-HCMV1/ARPE19-HCMV2). Null distributions were generated independently at each time-point, since each time-point was generated from an independent series of tandem-MS runs; and therefore the noise parameters were expected to vary.

Proteins with p-sample > 0.03 and a p-ratio > 0.05 were considered statistically significant. To identify co-regulated proteins across MRC5 and ARPE-19 cells, each protein was assigned to one of eight classes based on their fold-change trends and p-values (e.g. “A19-up/M5-up”, “A19-no-change/M5-up”, “A19-down/M5-down”, etc.). Enrichment analysis was performed using the Cytoscape^44^ plugin Cluego^46^.

### Data Availability

Raw MS-data are available through the ProteomeXchange Database (https://www.proteomexchange.org/). Processing scripts are available at Github (insert link).

---

**Figure S1.**
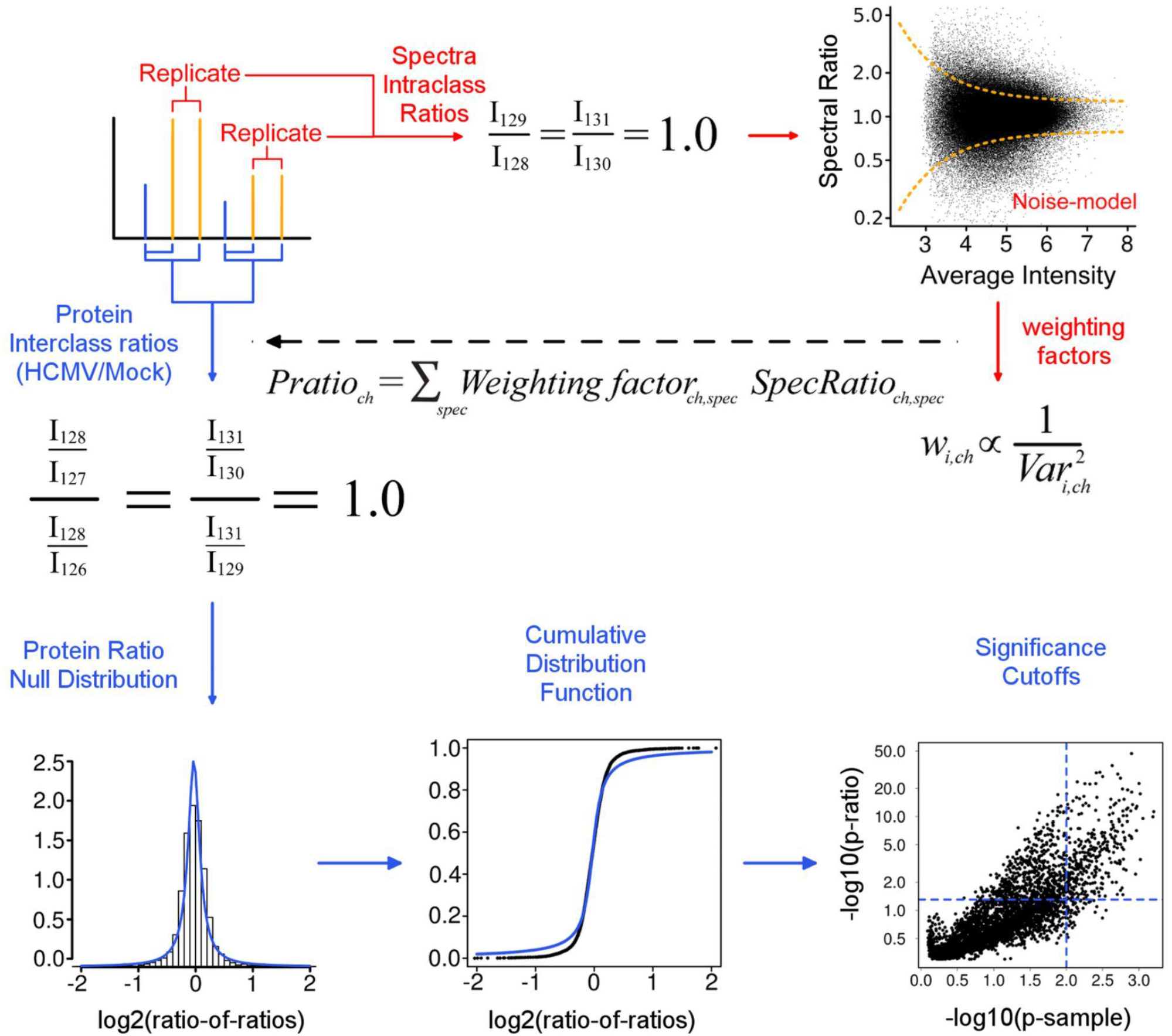
Mass-spectrometry Data Modeling using Isobar. A ratio noise-model yields weighting factors which are used to average reporter ion intensities from all spectra and peptides of the same protein. A null distribution is then constructed from the averaged protein ratios and used to identify significantly changed proteins. The ratio p-value is a measure of the ratio “quality”, loosely a metric of ratio signal-to-noise, accounting for the heteroscedasticity (intensity dependent noise) in the distribution of ratios. P-sample is probability that a given protein ratio is significantly different than the distribution of same-same, or “intraclass”, ratios. See Breitwieser, et al. (2011). Journal of Proteome Research 10, 2758–2766 for further details.

**Figure S2.**
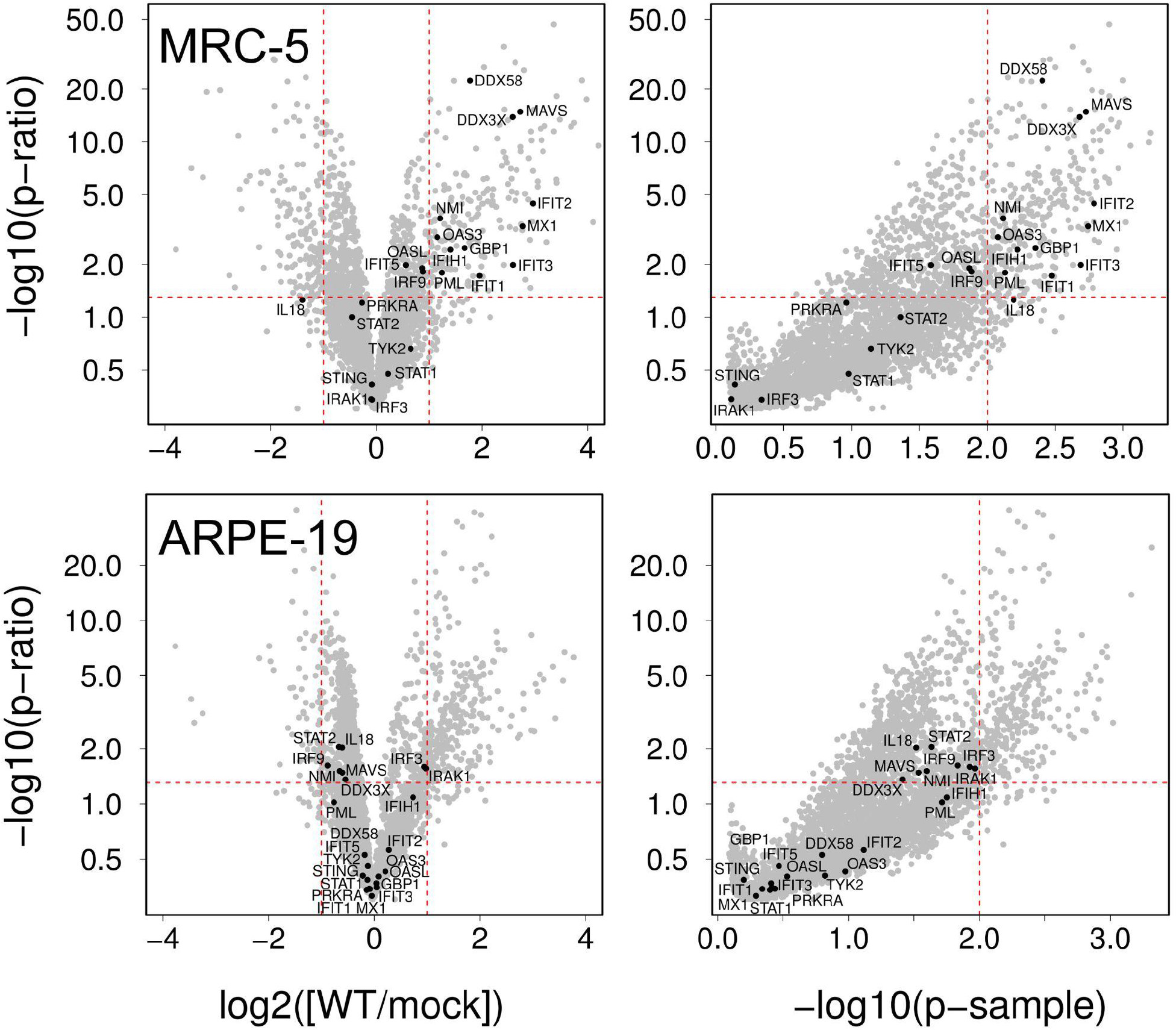

